# Intertumoral Genetic Heterogeneity Generates Distinct Tumor Microenvironments in a Novel Murine Synchronous Melanoma Model

**DOI:** 10.1101/2020.08.18.216259

**Authors:** Shuyang S. Qin, Booyeon J. Han, Alyssa Williams, Katherine M. Jackson, Rachel Jewell, Alexander C. Chacon, Edith M. Lord, David C. Linehan, Minsoo Kim, Alexandre Reuben, Scott A. Gerber, Peter A. Prieto

## Abstract

Synchronous metastatic melanoma, clinically defined as multiple lesions diagnosed within 6 months, has a poor prognosis. Despite recent advances in systemic immunotherapy, a majority of patients fail to respond or exhibit lesion-specific responses. While intertumoral heterogeneity has been clinically associated with lesion-specific therapeutic responses, no clear mechanism has been identified, largely due to the scarcity of preclinical models. We developed a novel murine synchronous melanoma model that recapitulates clinical intertumoral heterogeneity. We show that genetic differences between tumors generate distinct tumor immune microenvironments (TIME). These TIMEs can independently upregulate PD-1/PD-L1 expression in response to ongoing anti-tumor immunity and the presence of interferon-gamma. The simultaneous presence of multiple tumors can additionally alter the TIME of each tumor. As such, our model provides a unique approach to investigate the effects of intertumoral heterogeneity on mechanisms of immunotherapy resistance.

## Introduction

Metastatic melanoma carries a poor prognosis, with a median survival of less than 6 months[1]. Despite the curative potential of immunotherapies targeting the programmed cell death (PD)-1 pathway, less than 40% of stage IV melanoma patients respond and even fewer achieve remission[2, 3]. This variability is further complicated in synchronous metastatic melanoma, where multiple lesions are diagnosed within 6 months. The vast majority of these patients develop therapeutic resistance over time or lesion-specific mixed responses, wherein some metastases respond and others progress[3, 4]. The current standard of care requires a single biopsy to dictate therapy selection, which may not be representative of all tumors. This selection bias may negatively impact quality of life as patients often suffer adverse events without therapeutic benefit. Hence, novel methods are needed to personalize therapeutic selection for synchronous metastatic patients to generate optimal responses and minimize adverse events.

Whole exome sequencing of different metastatic lesions within the same patient demonstrates that melanoma metastases may share as little as 21% of somatic mutations[4]. Malignant melanoma has been identified as having one of the highest somatic mutational burdens, with a significant number of mutations being lesion-specific[5]. Intertumoral genetic heterogeneity has been clinically correlated to tumor immune microenvironmental (TIME) differences and subsequent lesion-specific therapeutic responses in multiple cancer types[4, 6]. Specifically, PD-1 immunotherapy response has been positively correlated to increasing tumor expression of PD-L1 (PD-1 ligand) and number of tumor-infiltrating CD8+ T cells[2, 7, 8]. Chronic PD-L1 expression is thought to upregulate of PD-1 on CD8+ T cells, inducing the T cell exhaustion program[9]. Exhausted T cells exhibit decreased proliferation and effector cytokine secretion, both of which could be potentially rescued by α-PD-1 immunotherapy[9–11]. One potent inducer of PD-L1 expression is interferon-gamma (IFN-γ), a critical cytokine in functional anti-tumor immune responses[12, 13]. The mechanisms underlying how these different components converge to determine the immunogenicity and immunotherapeutic response in synchronous metastatic melanoma are not well understood.

A major obstacle to studying the impact of intertumoral heterogeneity on anti-melanoma immunity is a scarcity of clinically relevant animal synchronous and metastatic models that recapitulate human disease. Approximately 50-60% of melanoma patients have the *BRAF*^*V600E*^ driver mutation, 60% have inactivating *CDKN2A* mutations and 5-20% have inactivating *PTEN* mutations[14]. The most commonly used mouse melanoma cell lines, including B16, harbor wildtype driver genes and thus cannot genetically represent the majority of human melanoma[15]. To remedy this problem, the Yale University Mouse Melanoma lines were developed with the YUMM 1.7 (YUMM) cell line containing the common *Braf*^*V600E/WT*^, *Pten*^−/−^, and *Cdkn2*^−/−^ driver mutations combination[16]. YUMMER 1.7 (YUMM<u>ER</u>) is a more immunogenic cell line derived from YUMM after multiple rounds of ultraviolet-B (UVB) irradiation in order to stimulate the most common mechanism for generating physiological mutations in melanoma[5, 17]. YUMM and YUMM<u>ER</u> cell lines share approximately 40% of somatic mutations[17]. Thus, these two cell lines are optimal candidates to model the observed intertumoral heterogeneity consistent with synchronous metastatic melanoma patients.

We developed a novel murine synchronous melanoma model using the YUMM and YUMM<u>ER</u> cell lines. We found that the YUMM and YUMM<u>ER</u> melanoma lines generate distinct TIMEs varying in tumor-infiltrating immune subtypes and surface marker expressions associated with T cell checkpoints. Furthermore, we discovered that ongoing inflammation, represented by the presence of IFN-γ, is critical for regional regulation of PD-1 and PD-L1 expression. Lastly, we demonstrated that the synchronous presence of multiple melanoma lesions alters TIMEs of individual tumors. Thus, we propose a new preclinical model of intertumoral heterogeneity for synchronous melanoma that may be studied to uncover mechanisms underlying lesion-specific immune and therapeutic responses.

## Results

### Expression of H2-K^b^ is increased in YUMM<u>ER</u> cells compared to YUMM cells in vitro

Whole exome sequencing revealed that the YUMM<u>ER</u> melanoma cell line contains roughly 60% of exonic mutations (1446 mutations) that are distinct from the parental YUMM line, with a significant portion secondary to UV-induced DNA damage[17]. We directly compared the two cell lines to determine if *de novo* somatic mutations in YUMM<u>ER</u> generated observable differences in RNA expression. Over 6000 (6197) genes were differentially expressed (**Fig. 1A**) between YUMM and YUMM<u>ER</u> *in vitro*. The YUMM<u>ER</u> cell line upregulated pathways associated with DNA damage repair and UV-induced stress, congruent with its generation by UVB-irradiation (**Fig. S1A-B**). Interestingly, YUMM<u>ER</u> also increased RNA expression of key genes associated with the induction of T cell exhaustion (**Fig. S1C**).

**Figure 1:**
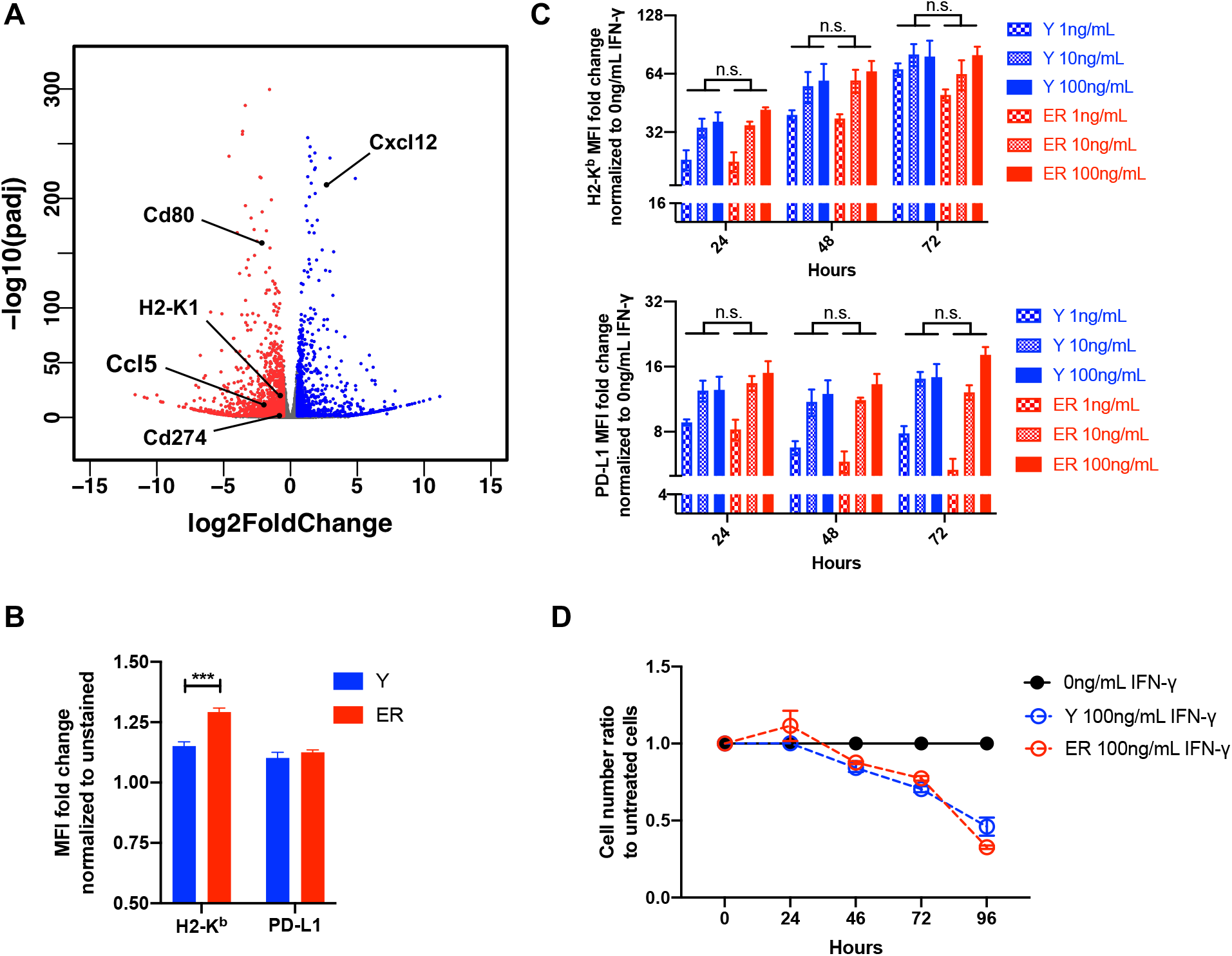
Y and ER differences *in vitro*. (A) Volcano plot highlighting differentially expressed genes in Y vs. ER cell lines. Colored genes are statistically significant with adjusted p-value < 0.05 and log 2-fold change > |0.5|. Genes of interest are labelled. (B) Normalized geometric mean fluorescence intensity (MFI) of baseline surface H2-K^b^ and PD-L1 on Y and ER cell lines. (C) Normalized geometric MFI of surface H2-K^b^ and PD-L1 on Y and ER cells cultured with exogenous IFN-γ over 72 hours. (D) Growth curves of cell number ratios of Y and ER cells with 100 ng/mL of exogenous IFN-γ compared to untreated. Data (mean ± SEM) in (B) – (D) are pooled, from 3 well/group/experiment, and representative of 2 independent experiments. ***p < 0.001, n.s. not significant.

As PD-L1 is an important inducer of T cell exhaustion[18], we analyzed its surface expression along with H2-K^b^, for which mRNA levels are elevated in YUMM<u>ER</u> cells. While YUMM and YUMM<u>ER</u> cell lines expressed similar levels of surface PD-L1 in culture, YUMM<u>ER</u> cells expressed higher basal H2-K^b^ levels (**Fig. 1B**). As the surface expression of both molecules may be induced by IFN-γ, we next assessed the responses of the two cell lines to exogenous IFN-γ. Both cell lines similarly upregulated H2-K^b^ and PD-L1 over time in response to varying IFN-γ dosages. (**Fig. 1C**, and **Fig. S2A-B**). Additionally, the cell lines exhibited similar growth kinetics with and without exogenous IFN-γ(**Fig. 1D** and **Fig. S2C**). Despite having no observable differences in the ability of YUMM and YUMM<u>ER</u> to upregulate surface H2-K^b^ and PD-L1 in response to IFN-γ in vitroγ, YUMM<u>ER</u> cells exhibited higher basal H2-K^b^ expression, suggesting that YUMM and YUMM<u>ER</u> tumors may elicit differential anti-tumor immunity *in vivo*.

### Genetic heterogeneity generates immunologically distinct microenvironments between synchronous YUMM and YUMM<u>ER</u> tumors

YUMM and YUMM<u>ER</u> melanoma cell lines contain similar proportions of shared and unique mutations as reported in previous analyses of synchronous metastatic melanomas, making them optimal cell lines to establish a murine model of synchronous melanoma[4, 17]. We therefore simultaneously injected YUMM and YUMM<u>ER</u> cells into the subcutaneous tissue of the left and right flanks of C57BL/6J mice (**Fig. 2A** and **Table S1**). Synchronous YUMM and YUMM<u>ER</u> tumors displayed similar growth kinetics *in vivo* with an initial period of equilibrium followed by tumor escape after day 20 (**Fig. 2B**). Although Luminex analysis of intratumoral cytokines/chemokines demonstrated that synchronous YUMM and YUMM<u>ER</u> tumors have similar levels of most cytokines and chemokines (**Fig. S3**), YUMM<u>ER</u> tumors exhibited elevated IFN-γ and CCL5, whereas contralateral YUMM tumors presented elevated CXCL12 (**Fig. 2C**).

**Figure 2:**
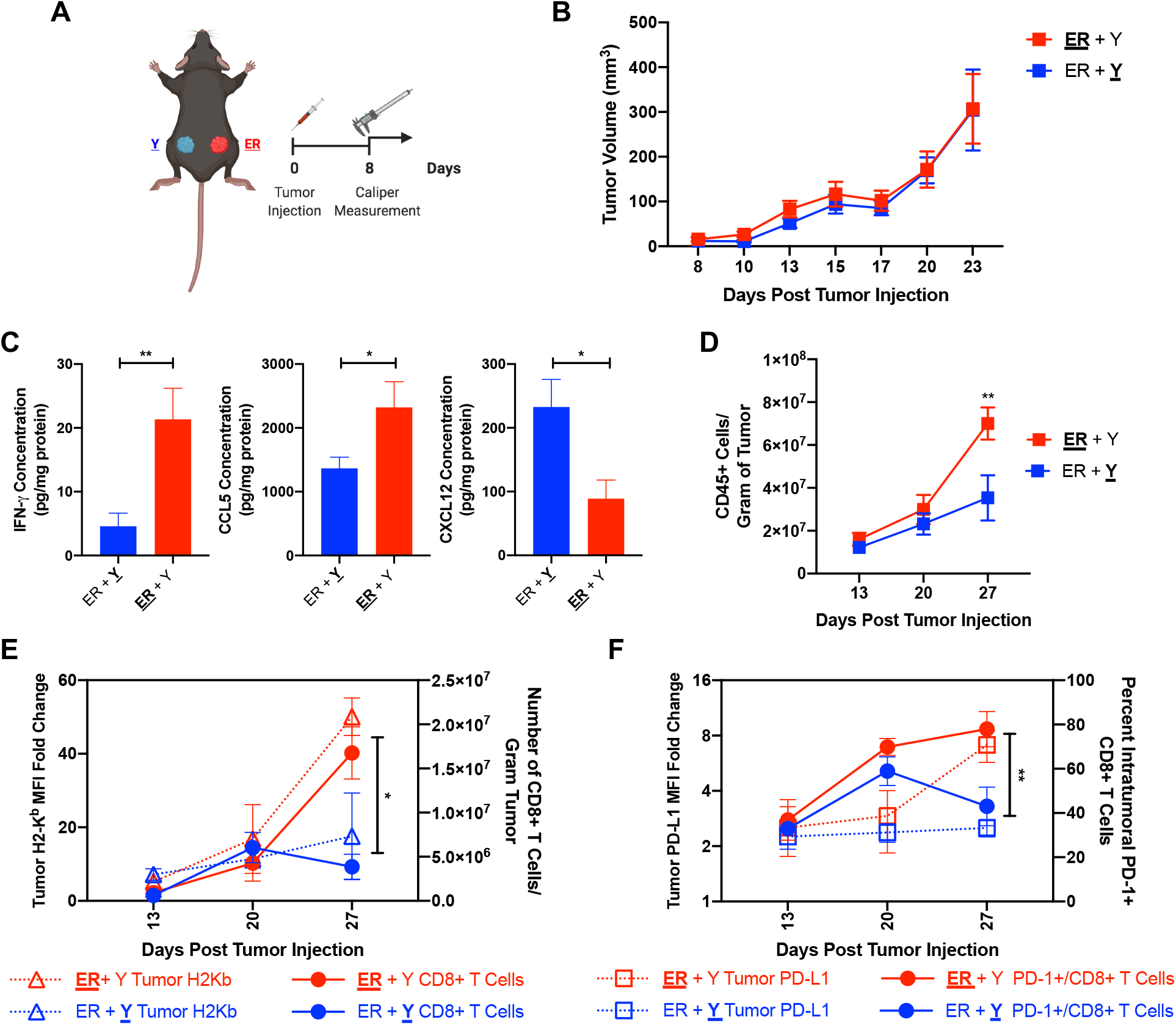
Synchronous Y and ER tumors establish distinct TIME *in vivo*. (A) Synchronous melanoma model schematic. The analyzed tumor in the synchronous model is **<u>underlined and bolded</u>** in subsequent figures. (B) Growth curves of individual Y (ER + **<u>Y</u>**) and ER (**<u>ER</u>**+ Y) tumors in synchronous melanoma mice. (C) Concentration of intratumoral IFN-γ, CCL5 and CXCL12 in synchronous Y or ER tumors isolated from synchronous melanoma mice on day 27. (D) Number of CD45+ immune cells over time in synchronous Y and ER tumors. (E) Relationship between tumor surface H2-K^b^ and recruited CD8+ T cells over time and (F) between tumor surface PD-L1 and percentage of PD-1+ CD8+ T cells over time in synchronous Y and ER tumors. Significance is indicated for each **<u>ER</u>**+ Y and ER + **<u>Y</u>** comparison per measured parameter. Data (mean ± SEM) in (B) – (F) are pooled, from 3-5 mice/group/experiment, and representative of at least 2 independent experiments. *p < 0.05, **p< 0.01.

We further analyzed the leukocyte recruitment into these tumors and observed increased CD45+ immune cell infiltration over time in synchronous YUMM<u>ER</u> tumors than in contralateral YUMM tumors (**Fig. 2D**). Interestingly, synchronous YUMM<u>ER</u> tumors significantly upregulated surface H2-K^b^ over time, accompanied by an increase in the number of tumor-infiltrating CD8+ T cells (**Fig. 2E**). At the same time, these YUMM<u>ER</u> tumors also upregulated surface PD-L1, corresponding to the accumulation of intratumoral PD-1+ CD8+ T cells (**Fig. 2F**). In contrast, contralateral YUMM tumors did not upregulate surface H2-K^b^ nor PD-L1, and were unable to recruit a high number of CD8+ T cells or maintain a high percentage of PD1+ CD8+ T cells over time (**Fig. 2E-F**). These results are congruent with the finding that synchronous YUMM<u>ER</u> tumors have elevated intratumoral IFN-γ compared to contralateral YUMM tumors. Our data suggest that the genetic differences between YUMM and YUMM<u>ER</u> cell lines are sufficient to establish different TIMEs on opposite flanks of the same mouse. Furthermore, these TIMEs may locally regulate the PD-1/PD-L1 checkpoint axis independently of one another.

### Interferon-gamma regulates tumor-specific PD-L1 upregulation and accumulation of PD-1+ CD8+ T cells

IFN-γ is a critical cytokine that shapes tumor development and has both pro-tumorigenic and anti-tumorigenic properties[13, 19]. Being one of the few differentially expressed cytokines in synchronous YUMM and YUMM<u>ER</u> tumors, we investigated the immunomodulatory effects of IFN-γ in our model. Synchronous YUMM and YUMM<u>ER</u> tumors grew faster and larger in *Ifn*g^−/−^ than wildtype mice (**Fig. 3A-B**). Further investigations into the individual TIMEs revealed that the lack of IFN-γ abolished the increase of CD45+ leukocytes observed in synchronous YUMM<u>ER</u> tumors, but had no effect on the number of immune cells in the YUMM tumors (**Fig. 3C**). The distribution of various immune subsets between *Ifng*^−/−^ synchronous YUMM and YUMM<u>ER</u> tumors were comparable and similar to that of wildtype synchronous YUMM tumors (**Fig. 3D**). In contrast, wildtype synchronous YUMM<u>ER</u> tumors exhibited increased CD8+ and macrophage/monocyte presence, suggesting a role for IFN-γ in preferential recruitment of these cell types into YUMM<u>ER</u> tumors (**Fig. 3E** and **Fig. S5**).

**Figure 3:**
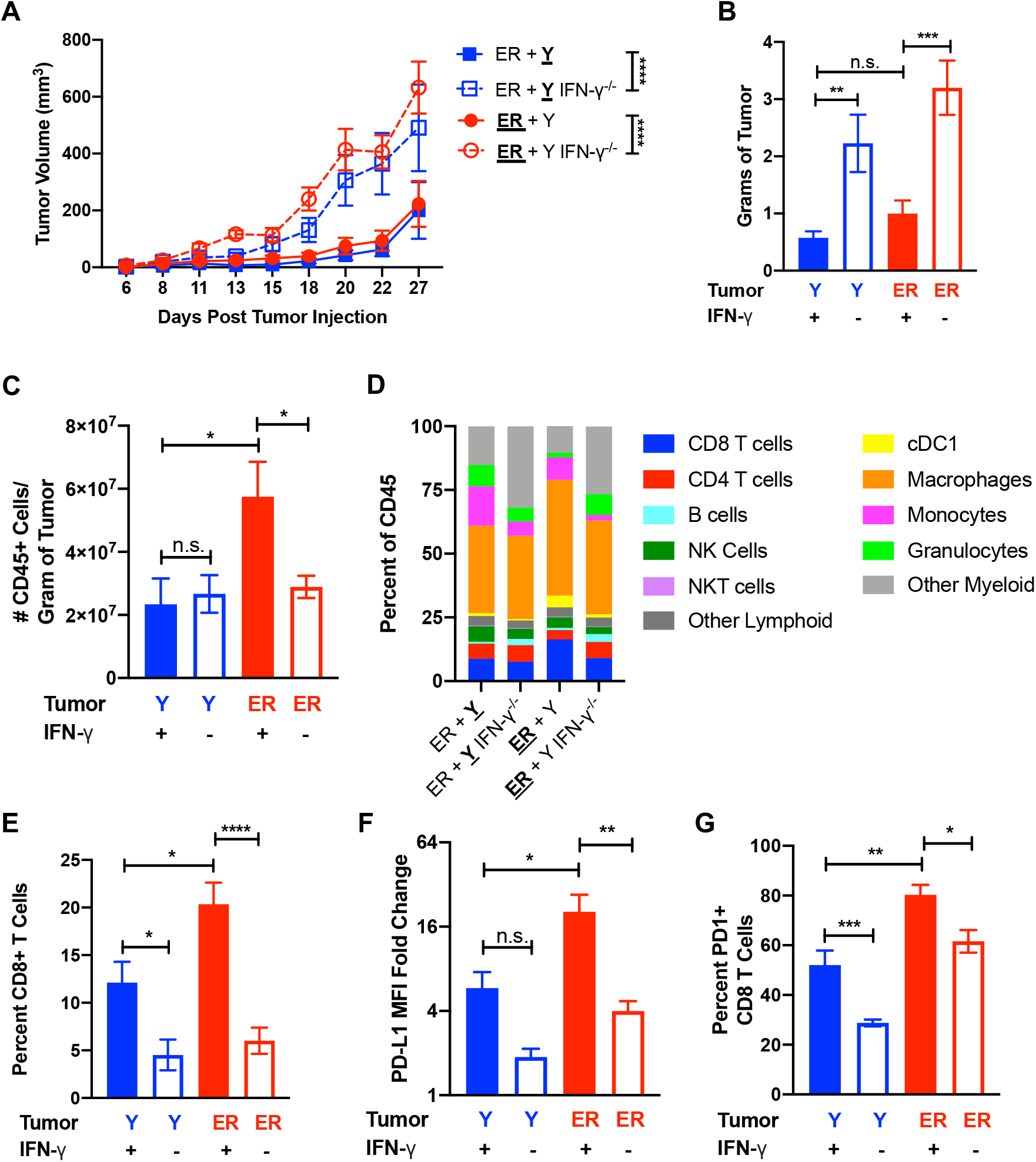
IFN-γ presence influences synchronous Y and ER growth and establishment of TIME. (A) Growth curves of individual Y (ER + **<u>Y</u>**) and ER (**<u>ER</u>**+ Y) tumors in synchronous melanoma model in wildtype and *Ifng*^−/−^ mice. (B) Tumor weight, (C) frequency of tumor-infiltrating CD45+ immune cells per gram of tumor, (D) overall immune subsets, and (E) percent of CD8+ T cells assessed on day 27. (F) Tumor surface PD-L1 and (G) percent of PD-1+ CD8+ T cells on day 27 of individual tumors in wildtype and *Ifng*^−/−^ synchronous mice. Data (mean ± SEM) in (B) are pooled, from 3-8 mice/group. Gating schematic for (D) shown in **Fig. S4.** Data in (C) – (G) are pooled from 3-5 mice/group/experiment and representative of at least two independent experiments. *p < 0.05, **p< 0.01, ***p < 0.001, **** p < 0.0001.

Since synchronous YUMM<u>ER</u> tumors can locally increase surface PD-L1 and accumulate more PD-1+ CD8+ T cells, we investigated the role of IFN-γ in regulating this pathway. YUMM<u>ER</u> tumors did not upregulate surface PD-L1 expression in *Ifng*^−/−^ mice (**Fig. 3F**). Interestingly, while the percentage of tumor-infiltrating PD-1+ CD8+ T cells decreased in both YUMM and YUMM<u>ER</u> tumors without IFN-γ, approximately 60% of YUMM<u>ER</u>-infiltrating CD8+ T cells were still PD-1+ in *Ifng*^−/−^ mice (**Fig. 3G**), implying the existence of another mechanism that allows these T cells to maintain high levels of surface PD-1 expression. Nevertheless, these data suggest that IFN-γ plays an important role in establishing the different TIMEs in synchronous melanoma tumors, especially in the local regulation of PD-1 and PD-L1 expression.

### Synchronous presence of genetically heterogeneous tumors alters anti-tumor immunity against individual tumors

To assess if tumors growing simultaneously in the same mouse can influence one another, we compared the synchronous YUMM and YUMM<u>ER</u> tumors against other tumor combinations (**Table S1**). While YUMM tumors exhibited the same growth kinetics in all contexts (**Fig. 4A**), synchronous YUMM<u>ER</u> growth was heavily influenced by the contralateral tumor (**Fig. 4B**). The presence of a contralateral YUMM tumor both facilitated synchronous YUMM<u>ER</u> tumor growth and prevented it from spontaneous rejection (**Fig. 4B-C**). This induction of YUMM<u>ER</u> tumor growth was highest with the genetically heterogeneous YUMM tumor, and not with an identical YUMM<u>ER</u> tumor nor with B16 tumor. Furthermore, this growth disparity disappeared in *Ifng*^−/−^ mice (**Fig. S6**), indicating that IFN-γ plays a role in this phenomenon.

**Figure 4:**
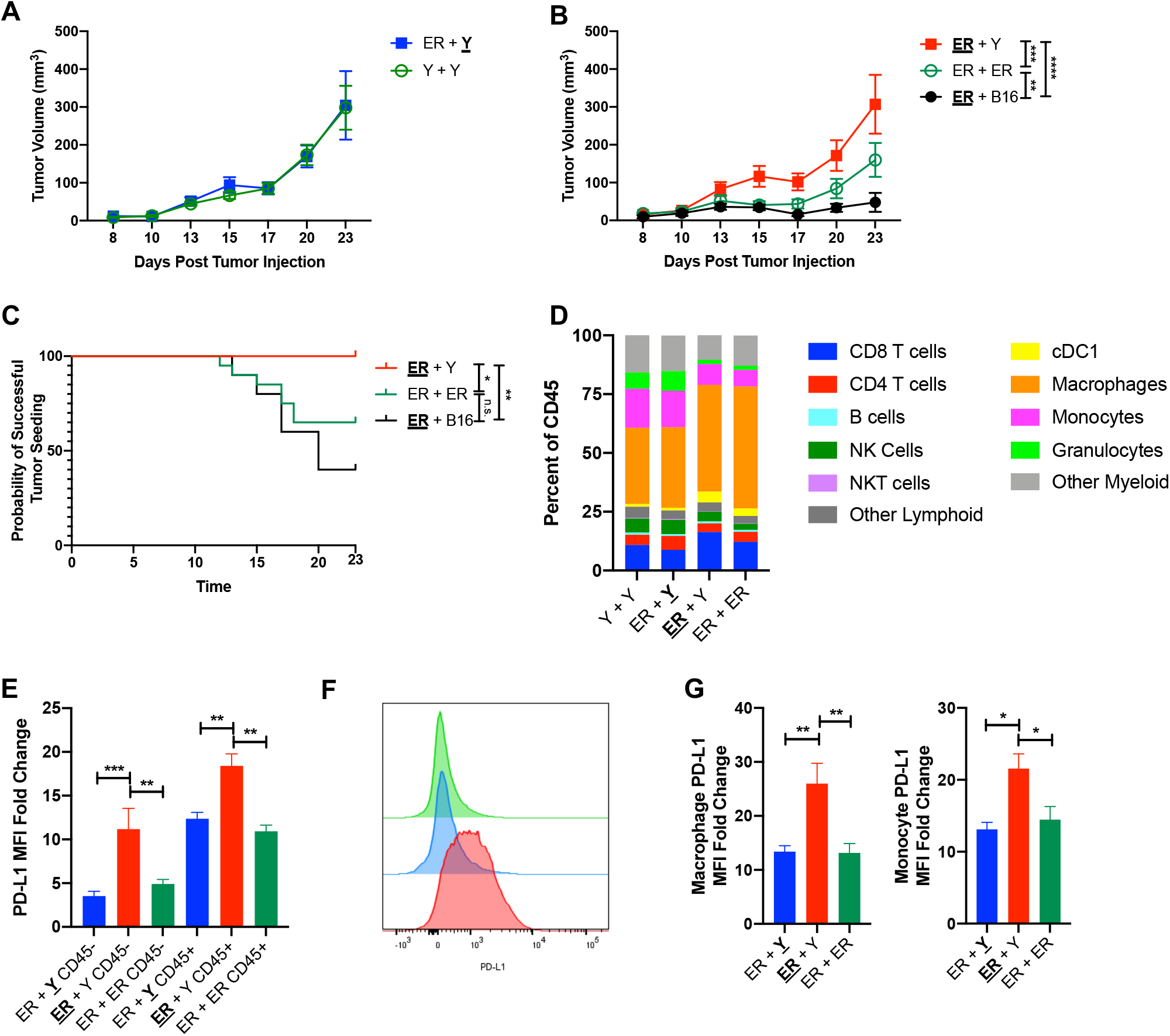
Synchronous Y + ER tumor combination alters ER TIME. (A) Growth curves of synchronous Y tumors from Y + ER and Y + Y mice. (B) Growth curves of synchronous ER tumors from ER + Y, ER + ER, and ER + B16 mice. (C) Percent of ER tumors successfully seeded over time. (D) Immune subset distribution and (E) normalized geometric MFI of tumor CD45- and CD45+ cell surface PD-L1 on individual synchronous tumors analyzed on day 27. (F) Representative plot and (G) summary of surface PD-L1 expression on tumor-infiltrating macrophages and monocytes day 27. Data (mean ± SEM) in (A) – (E) are pooled from 3-5 mice/group/experiment and representative of at least two independent experiments. Gating schematic for (D) shown in **Fig. S4.** Data in (F) and (G) are pooled from 3-5 mice/group. *p < 0.05, **p< 0.01, ***p < 0.001, **** p < 0.0001, n.s. not significant. Refer to Table S1 for tumor legend.

Dissection into the TIME differences of synchronous YUMM<u>ER</u> + YUMM tumors against other YUMM<u>ER</u> combinations demonstrated that the presence of contralateral YUMM only leads to minute differences in tumor-infiltrating immune populations (**Fig. 4D**). However, only these YUMM<u>ER</u> tumors upregulated surface PD-L1 expression on tumor cells and CD45+ tumor-infiltrating immune cells (**Fig. 4E**). This upregulation of PD-L1 on immune cells was largely contributed by a population shift in macrophages and monocytes in an IFN-γ dependent manner (**Fig. S7** and **Fig. 4F**) and was specific to the YUMM<u>ER</u> tumors in synchronous YUMM<u>ER</u> + YUMM mice (**Fig. 4G**). Overall, these data suggest that synchronous presence of genetically dissimilar YUMM and YUMM<u>ER</u> tumors can alter the YUMM<u>ER</u> tumor growth and TIME such that synchronous YUMM<u>ER</u> tumors selectively upregulate PD-L1 expression on tumor cells, tumor-infiltrating macrophages and monocytes as a potential mechanism of immunosuppression.

## Discussion

Genomic instability is one of the main mechanisms behind the generation of intertumoral genetic heterogeneity[20]. UV-induced DNA damage accounts for the majority of somatic mutations in malignant melanoma, making it the cancer with the highest mutational burden[5]. Studies have identified lesion-specific mutations between melanoma metastases within the same patient and correlated this intertumoral difference to heterogeneous TIME or mixed-response to systemic treatments[4]. However, the underlying mechanism has not yet been identified. Using our preclinical model of synchronous melanoma, we established that genetically dissimilar YUMM and YUMM<u>ER</u> melanoma cell lines generate distinct TIMEs of varying capabilities to elicit and affect tumor-infiltrating CD8+ T cells. The frequency and status of these CD8+ T cells ultimately determine lesion-specific anti-tumor immunity and immunotherapy response. Optimal combination therapy relies on selection of agents that cover all immunosuppressive mechanisms present in synchronous melanoma metastases.

While genetic differences may establish a predilection towards leukocyte recruitment, it is the manner by which tumors respond to ongoing inflammation and specifically the presence of IFN-γ, that determines their immunogenicity. YUMM tumors potentially evade the immune system by increasing CXCL12, whose high expression is associated with chemo-repulsion of T cells and exclusion of other CD45+ leukocyte infiltration[21]. In contrast, YUMM<u>ER</u> tumors upregulate IFN-γ, a critical cytokine in anti-tumor responses, and CCL5, a chemokine shown to recruit cross-presenting cDC1s into tumors[13, 22]. The presence of cDC1s is critical for T cell priming and migration as well as for spontaneous tumor rejection[23]. These YUMM<u>ER</u> TIMEs also contain significantly higher numbers of intratumoral CD8+ T cells. Despite these indicators of anti-tumor response, synchronous YUMM<u>ER</u> tumors from YUMM<u>ER</u> + YUMM mice display increased tumor growth compared to those from YUMM<u>ER</u> + YUMM<u>ER</u> mice. Additionally, synchronous YUMM<u>ER</u> tumors from YUMM<u>ER</u> + YUMM mice selectively upregulate surface PD-L1 on tumor and on tumor-infiltrating monocytes/macrophages in an IFN-γ dependent manner. This upregulation of PD-L1 is accompanied by increasing expression of PD-1 on intratumoral CD8+ T cells, a potential biomarker indicative of T cell exhaustion and inability to secrete effector cytokines [18, 24]. Thus, our data suggest the existence of additional immunosuppressive mechanisms, such as those involving the PD-1/PD-L1 axis, that are unique to synchronous melanoma. Detailed dissection of each individual tumor microenvironment is necessary to overcome therapeutic resistance. As such, assessment of individual melanoma metastases may be necessary to identify additional mechanisms of immunosuppression for optimal therapy selection and the prevention of lesion-specific therapeutic resistance.

A challenge with studying underlying mechanisms of tumor immunosuppression and immune evasion in metastatic melanoma is the lack of preclinical models that echo human melanoma. Here, we present a novel murine synchronous melanoma model using the YUMM and YUMM<u>ER</u> cell lines that recapitulate the genetic characteristics observed in human disease[4]. Using our model, we show that genetically dissimilar cell lines generate distinct TIMEs when synchronously implanted into opposing flanks of the same mouse despite displaying few appreciable phenotypic differences *in vitro*. The immunomodulatory capabilities of these TIMEs, including the regulation of PD-1 and PD-L1 checkpoint molecules, is dependent on the ability of the tumor to respond to local IFN-γ. Furthermore, we have evidence to suggest that the simultaneous presence of YUMM tumors can alter the contralateral YUMM<u>ER</u> TIMEs to facilitate immune escape.

Our novel murine synchronous melanoma model demonstrated that intertumoral genetic heterogeneity generates distinct TIMEs containing different immune escape mechanisms and that the presence of multiple, genetically similar lesions may alter the immune response of individual tumors. Further investigation is needed to determine the mechanism by which immunologically cold tumors facilitate growth of and suppress anti-tumor immunity against distal tumors. Nevertheless, this preclinical model will serve as a valuable tool to study the underlying mechanisms linking tumor genetics to tumor immunology *in vivo* and to identify potential targets to overcome lesion-specific therapeutic resistance secondary to intertumoral heterogeneity in synchronous metastatic melanoma. Based on our data potential targets include CCL5 and CXCL12. Initial data by others[25] suggest chemotactic inhibition may help augment immunotherapy by regulating trafficking at the level of the TIME. Further clinical trials based on this and our pre-clinical data are needed to further characterize this relationship.

## Methods

### *In vivo* Animal Studies

All *in vivo* procedures were performed in accordance with University of Rochester’s University Committee on Animal Resources approved guidelines. Six-to eight-week-old wildtype and age-matched *Ifng*^−/−^ (B6.129S7-*Ifng*^*tm1Ts*^/J) C57BL/6J mice were obtained from The Jackson Laboratory and from a generous gift by Edith Lord, PhD. Animals were given at least one week to acclimate before establishment of subcutaneous tumors.

### Cell Cultures

YUMM 1.7 cell line was purchased from ATCC. YUMM<u>ER</u> 1.7 cell line was generously gifted by Dr. Marcus Bosenberg. Cell cultures were maintained in DMEM:F12 media (Gibco) supplemented with 10% fetal bovine serum (Gibco), 1% penicillin/streptomycin (ThermoFisher Scientific) and 1% MEM non-essential amino acid solution (Gibco) at 37°C and 5% CO_2_.

### *In Vitro* Interferon-γ Studies

YUMM1.7 and YUMM<u>ER</u> 1.7 cell lines were plated in 6-well dishes and cultured in PBS (Gibco), 1ng/mL, 10ng/mL, or 100ng/mL of mouse IFN-γ (R&D Systems) in regular culture media. All cells within the well were collected at given timepoints to either be counted for total live cell numbers following trypan blue staining or resuspended in staining buffer for surface staining and flow cytometry analysis.

### Tumor Model and Tumor Volume Measurements

One × 10^6^ YUMM 1.7 or YUMMER 1.7 cells was simultaneously injected subcutaneously into opposing flanks of C57BL/6J mice in 100μL of PBS. Cell lines were detached with 0.25% trypsin/EDTA (Gibco) and resuspended in PBS (Gibco) for injection. Tumor growth was assessed with caliper measurements. Tumor volume was calculated by the formula 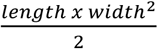.

### Tumor Single-Cell Suspensions

Mice tumors were individually excised, mechanistically dissociated, and digested in enzyme solution containing 10mM HEPES (Gibco), 1mg/mL Type IV Collagenase (Sigma-Aldrich), 150U/mL Type IV DNase I (Sigma-Aldrich) and 2.5U/mL Type V Hyaluronidase (Sigma-Aldrich) in RPMI (Gibco). Enzymatic digests were homogenized in the gentleMACS C tubes by alternating three times between 30-second pulse dissociation with gentleMACS dissociator and 10-minute incubation at 37°C. Homogenates were passed through 70μm filters and cells were resuspended in staining buffer (1mg/mL sodium azide and 10mg/mL BSA in PBS) to a final concentration of approximately 1-2 x10^6^ cells/100μL.

### Flow Cytometry

The following conjugated antibodies were used for staining: PerCP/Cy5.5 anti-mouse CD45 (30-F11, BD Pharmingen), FITC anti-mouse F4/80 (CI:A3-1, Abcam), APC/Cy7 anti-mouse CD8a (53.67, Lifescience), APC/Cy5.5 anti-mouse CD4 (GK1.5, Southern Biotechnology), APC anti-mouse PD-L1 (10F.9G2, Biolegend), BV786 anti-mouse CD11b (M1/70, Biolegend), BV711 anti-mouse CD103 (M290, BD Horizon), BV605 anti-mouse CD19 (1D3, Biolegend), PB anti-mouse Ly6G (1A8, Biolegend), PE/Cy7 anti-mouse Ly6C (HK1.4, Biolegend), PE/Cy5 anti-mouse IA/IE (M5/114.15.2, Invitrogen), PE/CF594 anti-mouse NK1.1 (PK136, BD Horizon), PE anti-mouse CD11c (N418, eBioscience), FITC anti-mouse CD106 (429, eBioscience), BV605 anti-mouse H-2Kb (AF6-88.5, BD Optibuild), PE anti-mouse CD119 (2E2, eBioscience), BV421 anti-mouse IA/IE (M5/114.15.2, BD Horizon), PE/Cy7 anti-mouse PD1 (RMP1-30, Biolegend), and PE/Cy5 anti-mouse CD3e (145-2C11, Biolegend). Cell surface antigens were stained for 30 minutes at 4°C in the dark. Following two staining buffer washes, the cells were fixed with BD Cytofix (BD Biosciences) for 20 minutes at 4°C in the dark before resuspension in staining buffer until analysis. Samples were run on a LSRII Fortessa (BD Biosciences). Fifty to one hundred thousand events were collected and analyzed using FlowJo software.

### Luminex Analyte Assay

Following sacrifice and excision, mouse tumors were homogenized with a tissue homogenizer in 700μL of Cell Lysis Buffer 2 (R&D Systems) containing 1x Halt Protease Inhibitor Cocktail (ThermoFisher Scientific). Tissues were lysed on ice for 30 minutes with gentle agitation. Magnetic Luminex Assays were performed with a Mouse Premixed Cytokine/Chemokine Multi-Analyte Kit (R&D Systems) per manufacturer’s instructions. Microplates were run on a Bio-Flex 200 system (Bio-Rad) collecting 50-100 beads per target with less than 20% aggregate. Pierce BCA Protein Assays (ThermoFisher Scientific) was performed on the remaining lysates following manufacturer’s instructions to determine total protein concentrations. Analyte concentrations were normalized to total protein concentration for each sample into pg analyte/mg protein.

### RNA Sequencing and Analysis

Cells were detached from the culture with 0.25% trypsin and lysed in RLT Plus buffer containing 1% β-mercaptoethanol. Lysates were homogenized with QIAShredder spin columns, and RNA was purified using the RNeasy Micro Kit (QIAGEN) following manufacturer’s instructions. RNA sequencing and preliminary differentially expressed gene analysis was performed by the University of Rochester Genomics Research Center. RNA quality was assessed using an Agilent Bioanalyzer (Agilent) and cDNA libraries were constructed with TruSeq RNA Sample Preparation Kit V2 (Illumina) according to manufacturer’s instructions. Sequencing was performed on HiSeqTM 2500 (Illumina). Raw reads were demultiplexed using bcl2fastq version 2.19.1 and mapped to the Mus musculus reference genome (GRCm38 + Gencode-M22 Annotation) using STAR_2.7. Differential expression analysis was performed using DESeq2-1.22.1 with a P-value threshold of 0.05, within R version 3.5.1. Subsequent pathway analysis was performed using Qiagen Ingenuity Pathway Analysis (IPA) software.

### Quantification and Statistical Analyses

Prism 8 software (GraphPad) was used for all statistical analyses with p-values < 0.05 determined statistically significant. Tumor growth data was analyzed by mix-model analysis with Tukey’s multiple comparisons test at each timepoint. Tumor rejection data was analyzed by the logrank test. All flow cytometry gating was performed using FlowJo 10 software (FlowJo) with one-way ANOVA or paired t-test analyses to assess for statistically significance differences in cell density or geometric mean intensity between various groups of tumors or cell lines. All diagrammatic figures were created with BioRender.

## Acknowledgements

Supported by grants from the NIH (R01CA230277 to S.A.G.; R01CA168863 to D.C.L, T32GM007356 training grant to S.S.Q) and University of Rochester Medical Center start-up fund to P.A.P. We thank the core facilities at the University of Rochester Medical Center; specifically, Dr. Timothy Bushnell in the Flow Cytometry Shared Resource Core, and Dr. Cameron Baker in the Genomics Research Core.

