## Supplemental Figures for "Intertumoral Genetic Heterogeneity Generates Distinct Tumor Microenvironments in a Novel Murine Synchronous Melanoma Model"

| Name | Tumor Analyzed | Tumor Combinations in Mice |
|---|---|---|
| **ER** + Y | YUMMER | **YUMMER** + YUMM |
| ER + **Y** | YUMM | YUMMER + **YUMM** |
| ER + ER | YUMMER | YUMMER + YUMMER |
| Y + Y | YUMM | YUMM + YUMM |
| **ER** + B16 | YUMMER | **YUMMER** + B16 |

**Table S1:** Tumor legend notation.

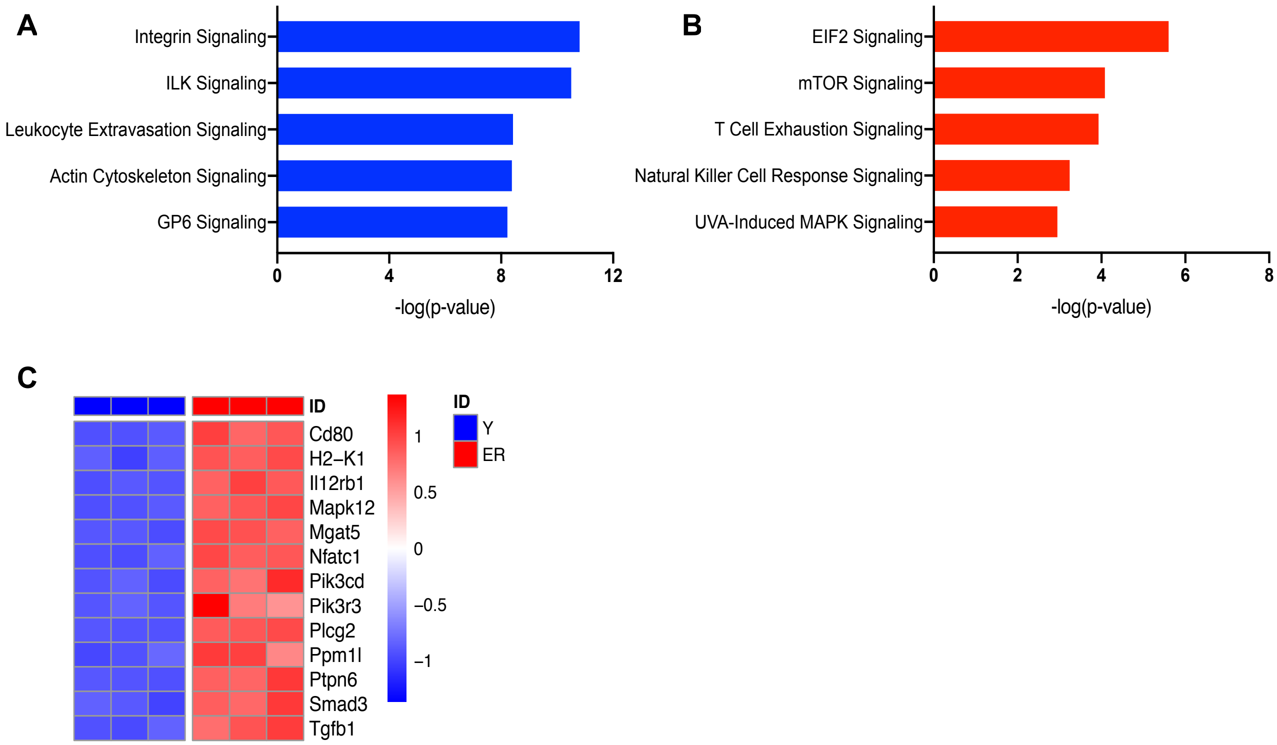


**Figure S1:** Top up-regulated pathways in Y and ER *in vitro*. Top 5 most upregulated pathways in Y (A) and ER (B) cell lines identified by IPA analysis. (C) Heatmap showing statistically significant differentially expressed genes involved in T cell exhaustion induction pathway according to IPA analysis.

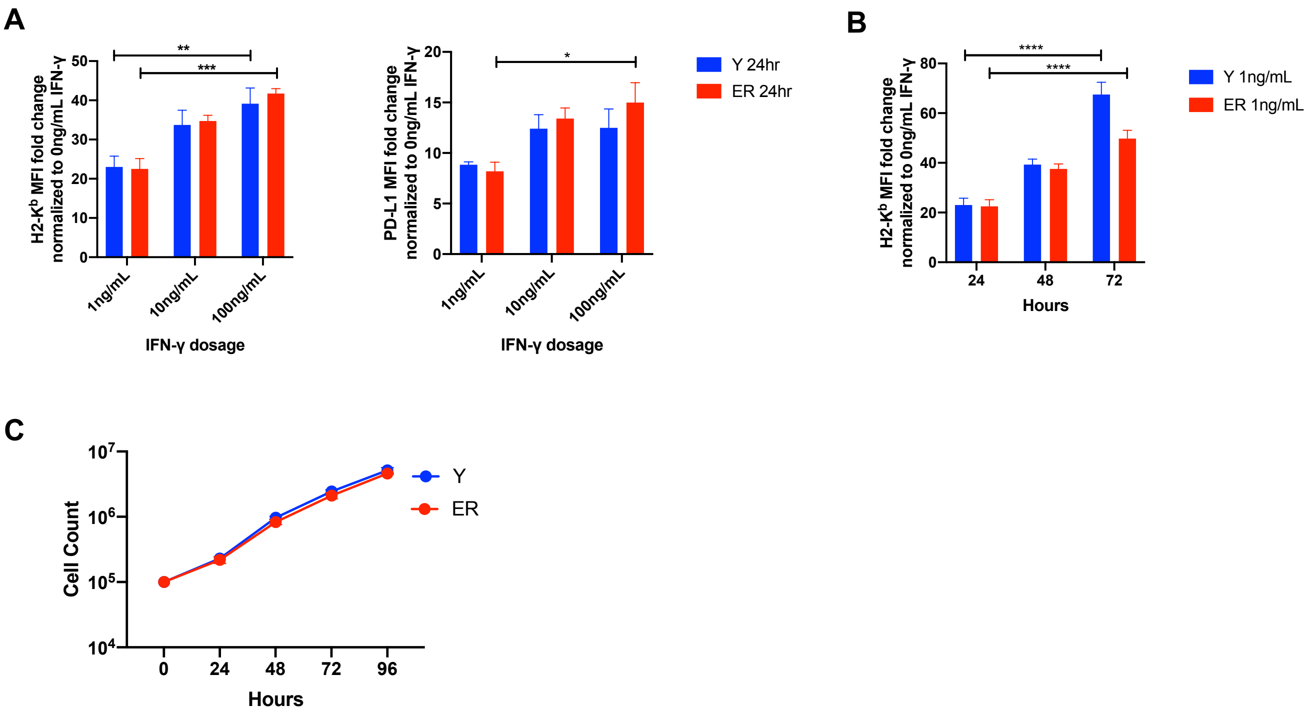


**Figure S2:** (A) Normalized geometric mean fluorescence intensity (MFI) of surface H2-K$^b$ and PD-L1 on Y and ER cell lines after 24 hour exposure to varying levels of exogenous IFN-γ. (B) Normalized geometric MFI of surface H2-K$^b$ on Y and ER cells cultured with 1ng/mL IFN-γ over time. (C) Growth curves of Y and ER without IFN-γ. Data (mean ± SEM) are pooled, from 3 well/group/experiment, and representative of 2 independent experiments. $*p < 0.05$, $**p < 0.01$, $***p < 0.001$, $****p < 0.0001$.

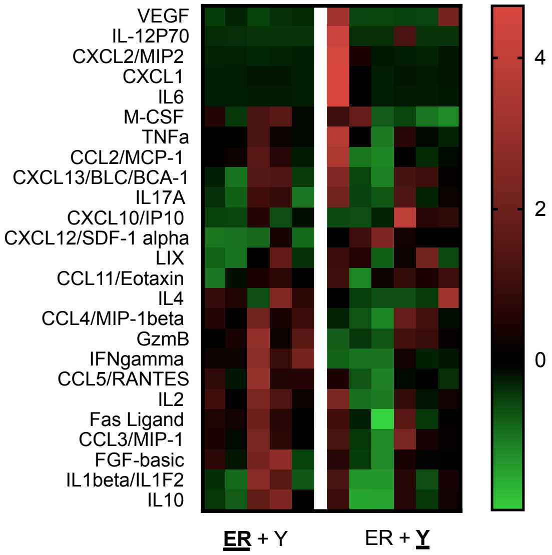


**Figure S3:** Relative concentrations of intratumoral cytokines/chemokines in synchronous Y (ER + **Y**) and ER (**ER** + Y) tumors.

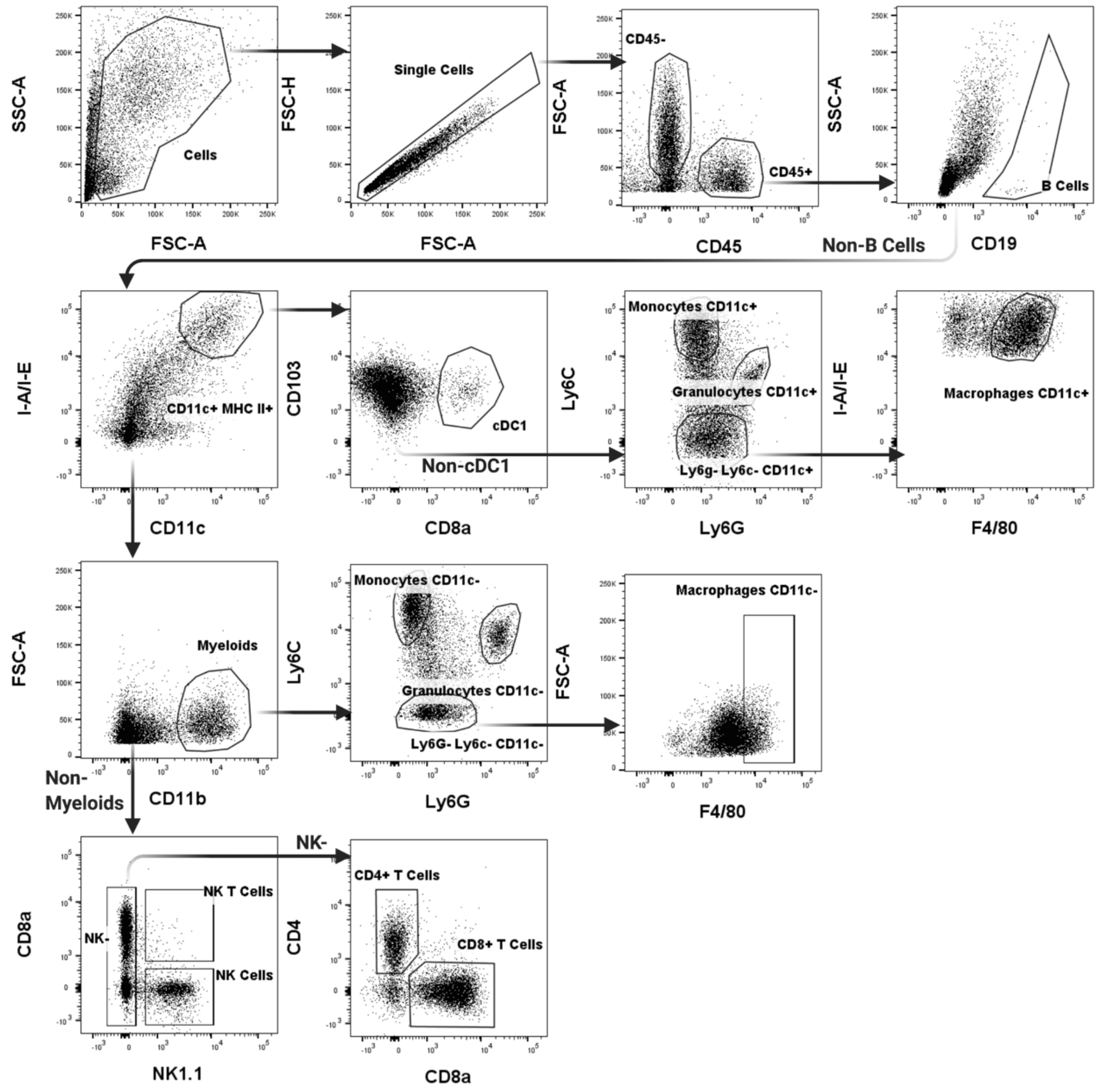


**Figure S4:** Flow cytometric tumor-infiltrating immune subset gating scheme. Total macrophages, monocytes and granulocytes refer to the concatenated sum of respective CD11c+ and CD11c- populations.

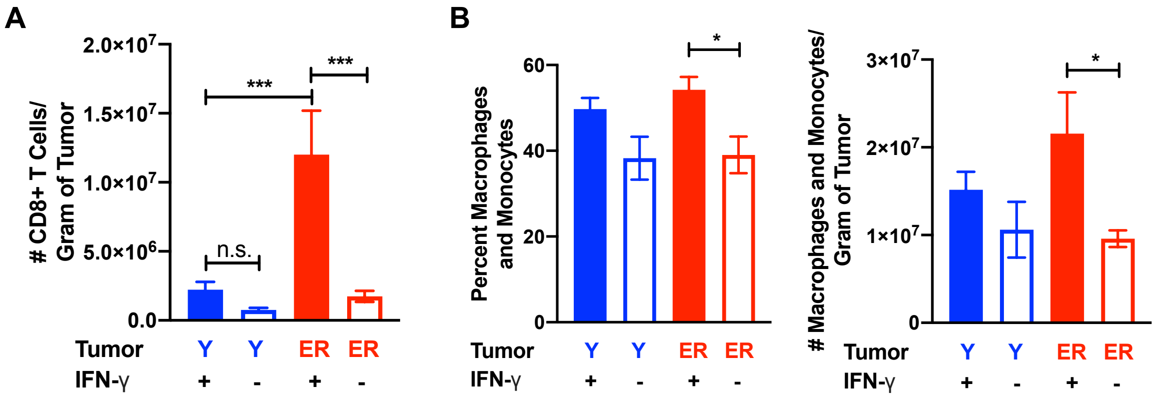


**Figure S5:** Frequency of tumor-infiltrating CD8+ T cells (A), percent and frequency of tumor-infiltrating macrophages/monocytes (B) in synchronous Y (ER + **Y**) and ER (**ER** + Y) tumors of wildtype and *Ifng*$^{-/-}$ mice on day 27. Data (mean ± SEM) are pooled from 3-5 mice/group/experiment and representative of at least two independent experiments *p < 0.05, *** p < 0.001.

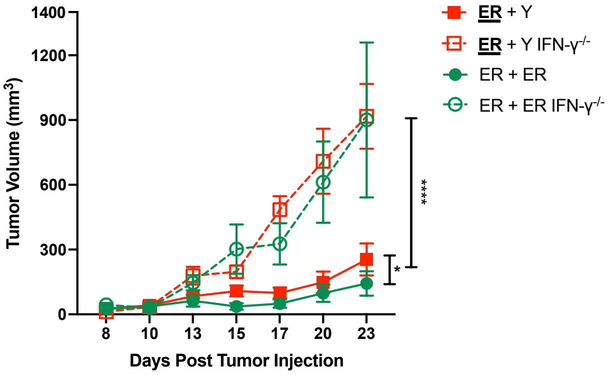


**Figure S6:** Growth curves of synchronous ER tumors from <u>ER</u> + Y and ER + ER mice in wildtype and *Ifng*$^{-/-}$ mice. Data (mean ± SEM) are pooled from 3-5 mice/group *p < 0.05, ****p < 0.0001.

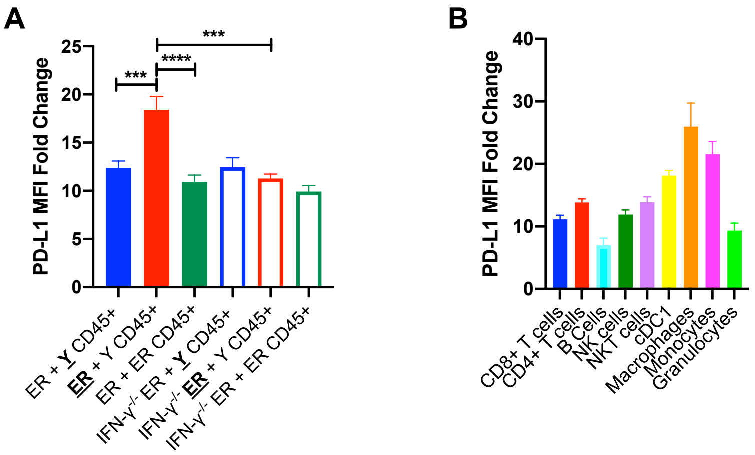


**Figure S7:** (A) CD45+ surface PD-L1 geometric MFI on day 27 of individual tumors in synchronous wildtype and *Ifng*$^{-/-}$ mice. (B) Surface PD-L1 expression measured by geometric MFI on different tumor-infiltrating immune subsets in ER + Y on day 27. Data (mean ± SEM) are pooled from 3-5 mice/experiment from two independent experiments. ***p < 0.001, **** p < 0.0001.
